# Gestational day 12 moderate prenatal alcohol exposure produces sex-specific social impairments and attenuates prelimbic excitability and amygdala-cortex modulation of adult social behavior

**DOI:** 10.1101/2022.05.02.490324

**Authors:** Kathryn R. Przybysz, Mary B. Spodnick, Julia M. Johnson, Elena I. Varlinskaya, Marvin R. Diaz

## Abstract

Lifelong social impairments are common in individuals with prenatal alcohol exposure (PAE), and preclinical studies have identified gestational day (G)12 as a vulnerable timepoint for producing social deficits following binge-level PAE. While moderate (m)PAE also produces social impairments, the long-term neuroadaptations underlying them are poorly understood. Activity of the projection from the basolateral amygdala to the prelimbic cortex (BLA→PL) leads to social avoidance, and the PL alone is implicated in negative social behaviors, making each of these potential candidates for the neuroadaptations underlying mPAE-induced social impairments. To examine this, we first established that G12 mPAE produced sex-specific social impairments lasting into adulthood. We then chemogenetically inhibited the BLA→PL using Clozapine N-Oxide (CNO) during adult social testing. This revealed that CNO reduced social investigation in control males, but had no effect on mPAE males or females of either exposure, indicating that mPAE attenuated the role of this projection in regulating male social behavior and highlighting one potential mechanism by which mPAE affects male social behavior more severely. Using whole-cell electrophysiology, we also examined mPAE-induced changes to PL pyramidal cell physiology and determined that mPAE reduced the excitability of these cells, likely due to increased suppression by inhibitory interneurons. Overall, this work identified two mPAE-induced neuroadaptations that last into adulthood and which may underlie the sexspecific vulnerability to mPAE-induced social impairments. Future research is necessary to expand upon how these circuits modulate both normal and pathological social behavior, and to identify sex-specific mechanisms leading to differential vulnerability in males and females.

## Introduction

Alcohol exposure during prenatal development produces many negative outcomes in exposed offspring, and ~5% of individuals in the United States are diagnosed with a Fetal Alcohol Spectrum Disorder (FASD) [1]. Mental illness is widespread among individuals with FASD, and mood-related disorders like anxiety make up ~68% of FASD-associated mental illness across the lifespan [2]. One of the most common behavioral symptoms of FASD is social impairment, including social anxiety, which often lasts into adulthood [3–10].

Preclinical models of prenatal alcohol exposure (PAE) have recapitulated human literature, demonstrating that exposure to chronic or binge-level ethanol during prenatal development disrupts social behavior in young (juvenile/adolescent) [11–16] and adult [11, 13–15] rodents. Specifically, an acute, extreme binge-level exposure reduced social investigation and social motivation across several ages depending on sex and the timing of exposure [11, 14, 15], while other social impairments such as reductions in social contact and play were age- and sex-specific [11, 13–15]. Throughout these studies, male social behavior was more severely affected by PAE while also being impacted earlier in life compared to females. Based on the gaps left by these preclinical studies, it is necessary to gain a better understanding of neural mechanisms underlying PAE-induced social impairments in males and females.

Epidemiological studies have demonstrated that pregnant women more often consume alcohol in the low-moderate range than the binge-level range, and that low-level consumption increases across pregnancy [17–21]. While fewer studies have examined the long-term effects of moderate PAE (mPAE), similar effects on social behavior have been observed [7] across development [22–31], and with males more affected than females [22, 23, 25]. We have demonstrated that a high-dose ethanol exposure on gestational day (G) 12 sex-specifically impairs social behavior across multiple rat strains [11, 14], while *m*PAE targeting G12 increases general anxiety-like behavior in adolescent males only [32]. G12 is a period of development during which brain structures involved in the expression and regulation of anxiety-related behaviors, such as the amygdala, are formed [33, 34].

Social interaction requires the recruitment of many different brain regions [35, 36], and the amygdala and the medial prefrontal cortex (mPFC) are two areas that are critical for social behaviors relating to emotional processing. The basolateral amygdala (BLA) and the prelimbic subdivision of the mPFC (PL) specifically are implicated in regulating social behavior, with activity of these subregions associated with negative social behaviors like social anxiety and social avoidance [37–46]. These subregions are also interconnected, and activity of the neural projections specifically from the BLA to the PL (BLA→PL) is associated with reduced social behavior in male mice [47], highlighting the role of this projection in specifically modulating negative social behaviors. Notably, both the BLA and the PL are functionally vulnerable to PAE, as BLA excitability is increased [48, 49], and mPFC excitability is decreased [50–52] after PAE. Increased inhibitory interneuron populations in different mPFC subregions, including the PL, support this shift in excitability [50, 52, 53]; however, PL function following PAE has not been directly assessed. Furthermore, given the role of the BLA→PL projection in social anxiety and social avoidance, improving our understanding of how the PL may be functionally altered by PAE to produce social impairments like those described above, has not been examined.

The objective of the present work was three-fold: 1) determine whether G12 *m*PAE would impact social behavior, like what we have found with G12 binge-level PAE, 2) investigate the role of the BLA→PL projection in regulating PAE-induced social impairments, and 3) assess G12 mPAE-induced neuroadaptations in PL function.

## Methods

Refer to Supplementary Materials for expanded methodology

### Subjects & Moderate Prenatal Alcohol Exposure

Male and female Sprague-Dawley rats bred in our colony were used for all experiments, and breeding and G12 mPAE procedures were conducted as described previously [32]. Adult male and female breeders were obtained (Envigo, Indianapolis, IN) and allowed to acclimate to the colony for at least one week prior to breeding. On G12, pregnant dams were placed into a vapor inhalation chamber for six hours (09:00-15:00) and exposed to either room air or vaporized ethanol. This exposure produces peak blood ethanol concentrations in the moderate range (75-100 mg/dl [32, 54]). The dams were allowed to complete their pregnancies, and pups were weaned on P21 and assigned to one of three experiments as described below. Early adolescent (postnatal day (P) 28), late adolescent (P42), and adult (P77) rats were used for initial social interaction testing, and adult (P70+) rats were used for social behavior testing with chemogenetic manipulation and for electrophysiology experiments. In all respects, maintenance and treatment of the animals were in accord with guidelines established by the National Institutes of Health, using protocols approved by the Binghamton University Institutional Animal Care and Use Committee.

### Social interaction test

Social interaction testing was conducted as previously described [11]. Male and female experimental rats were placed into a two-compartment testing chamber for a 30-minute habituation period, followed by a ten-minute interaction period with an unfamiliar social partner of the same age and sex. All behaviors were video recorded and saved for later scoring. Behaviors scored included frequencies of social investigation (sniffing of the partner), contact behavior (grooming of and climbing over/under the partner), and play behavior (pouncing, following/chasing, pinning). Social motivation via social preference/avoidance was measured as crossovers from one compartment of the testing apparatus toward (preference) or away from (avoidance) the partner, and a social preference/avoidance coefficient was calculated [coefficient (%)=(crossovers toward partner–crossovers away from the partner)/(total crossovers toward/away)x100].

### Viral infusion and cannulation surgery

In a different subset of animals, offspring underwent a viral infusion/cannulation surgical procedure between P70-80. Rats were induced with 3-5% isoflurane anesthesia and maintained on 2-4% throughout surgery. Craniotomies were made above the BLA and PL, and rats received an infusion of either an inhibitory chemogenetic virus (pAAV-CaMKIIa-hM4D(Gi)-mCherry) or a control virus (pAAV-CaMKIIa-mCherry) bilaterally into the BLA (0.1 μL/min, 0.5 μL/side; coordinates: AP: +2.5, ML: ±5.0, and DV: −8.8 mm relative to Bregma) using a microsyringe pump (Stoelting). Each rat was then implanted with a bilateral guide cannula (26-gauge, 1 mm between guides; Plastics One, Roanoke, VA) such that the cannula terminated 1 mm above the PL (coordinates: AP: +3.5, and DV: −2.5 relative to Bregma), and a dummy cannula and dust cap were placed to maintain the patency of the cannulae until testing. Postoperative care was conducted once per day for 10 days after surgery, and twice per week thereafter until testing.

### Microinjection procedure

6 weeks after surgery, all rats underwent the social interaction testing procedure as described above. Immediately before testing, each rat received a microinjection of either Clozapine N-Oxide (CNO, 300 μM, 0.5 μL per side) or vehicle (artificial cerebrospinal fluid, ACSF). Each rat was tested twice 48 hours apart, each day receiving either CNO or ACSF with the order of drugs counterbalanced across groups. After the second day of testing, rats were transcardially perfused with 4% paraformaldehyde and brains were preserved for histological analysis. Viral infusion sites were confirmed via fluorescence microscopy using immunofluorescence to visualize mCherry in coronal BLA slices, and cannula placements were confirmed by preparing coronal mPFC slices from fixed tissue.

### Whole-cell electrophysiology

All electrophysiological procedures were conducted in adult (P80+) offspring. Rats were anesthetized with 4-5% isoflurane and brains were removed and submerged in ice-cold NMDG cutting solution. 350 μm PL-containing slices were made using a vibratome (Leica Microsystems), and allowed to incubate in artificial cerebral spinal fluid (ACSF) at 34°C for at least 40 min prior to experiments. All experiments were conducted within 4 hours of slice preparation.

Electrophysiological procedures were conducted as previously described [54–56]. Recordings were made from PL layer 2/3 (PL2/3) pyramidal cells, whose identity was initially based on morphology and membrane properties and subsequently confirmed based on the resting membrane potential (RMP) and pattern of action potential (AP) firing [57, 58].

Intrinsic excitability of pyramidal cells was assessed at RMP in a bath containing only ACSF. After obtaining an RMP reading for each cell in zero-current configuration, a series of depolarizing current steps (increasing steps of 10 pA beginning at −100 pA, 500 ms step duration) was applied to elicit APs in current clamp configuration and the following parameters were assessed: rheobase, membrane potential of the first AP, time to first AP, AP peak, and the number of AP fired with increasing current steps. To assess the role of GABAergic inhibition in modulating pyramidal cell firing, identical experiments were conducted in a separate subset of neurons. A first recording was made of the basal firing in ACSF alone, then a second identical recording was conducted in the same cell while pharmacologically blocking GABAA receptors with gabazine (10 μM). Following intrinsic excitability recordings, excitatory postsynaptic currents (EPSCs) were recorded in voltage-clamp configuration. Baseline spontaneous (s)EPSCs were recorded during the continued application of gabazine, followed by miniature (m)EPSC recordings in the presence of tetrodotoxin (0.5 μM) to assess AP-independent transmission.

### Data analysis

All data was done by a researcher blind to the experimental condition. All statistical analyses, including *t*-tests, between-subjects ANOVAs, and mixed ANOVAs were conducted using Prism 6 (GraphPad), and data from males and females were analyzed separately [59–61].Following significant main effects or interaction, Bonferroni corrected post-hoc tests were conducted to detect significant differences between groups. The significance threshold was set at p < 0.05, and data are represented as mean ± SEM.

## Results

### G12 mPAE produces sex-specific social impairments lasting into adulthood

We first examined the social behavior of early adolescent, late adolescent, and adult rats who had been exposed to G12 mPAE. In males, the ANOVA revealed a main effect of G12 exposure on social investigation [*F*(1,46)=10.12, *p*=0.0026](*Fig. 1A*), social motivation [*F*(1,46 =15.16, *p*=0.0003](*Fig. 1B*), and social contact behavior [*F*(1,49)=7.68, *p*=0.0079] (*Fig. 1C*) with mPAE males exhibiting reductions compared to air-exposed males. There was no main effect of age or age x exposure interaction for these behaviors. The findings from these three variables indicate that mPAE males exhibited significant social impairment lasting into adulthood. We also found a main effect of age on social play [*F*(2, 46)=6.79, *p*=0.003], indicating that social play decreased across ontogeny [adults vs early adolescents: *t*(1,46)=3.56, *p*=0.0056; adults vs. late adolescents: *t*(1, 46)=2.55, *p*=0.0422](*Fig. 1D*), but no effect of exposure or an age x exposure interaction.

**Figure 1.**
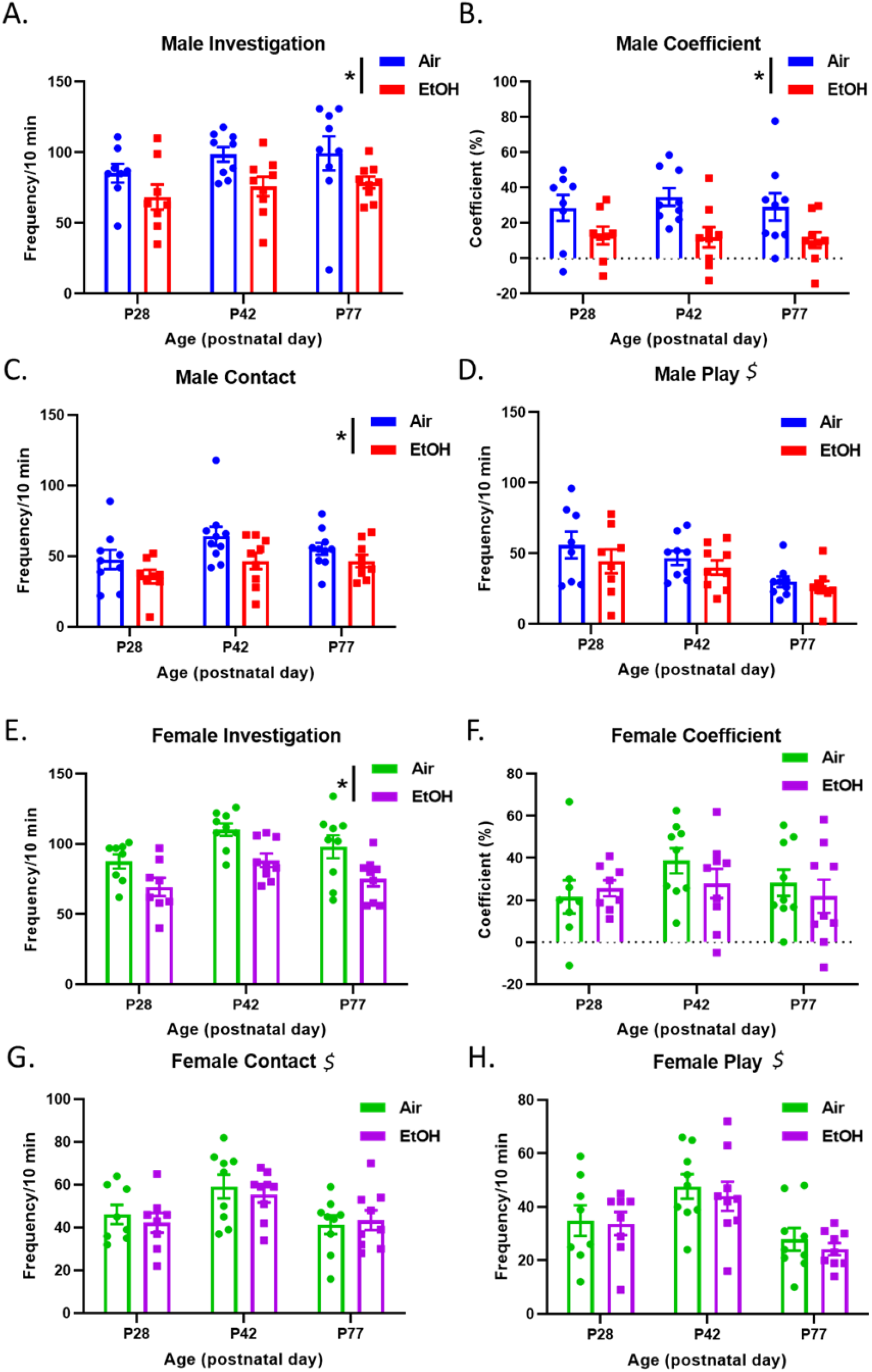
The impact of G12 mPAE on social behaviors of male and female rats tested as early adolescents (P28), late adolescents (P42) or adults (P77). G12 mPAE males had reduced **(A)** social investigation, **(B)** social preference, **(C)** and social contact, but not **(D)** social play. For social play, males showed a decrease in play across age, regardless of mPAE exposure. G12 mPAE females had reduced **(E)** social investigation but not **(F)** social preference, **(G)** social contact, or **(H)** social play. Females also exhibited age-specific changes of **(G)** social contact and **(H)** social play, with late adolescents showing the highest levels of social contact and play. * indicates a significant effect of exposure (*p*<0.05), $ indicates a significant effect of age (*p*<0.05).

Female mPAE offspring also showed reduced social investigation, with a main effect of G12 exposure [*F*(1,46)=19.28, *p*<0.0001]. We also found a main effect of age [*F*(2, 46)=6.376, *p*=0.0036], but no age x exposure interaction [*p*=0.91](*Fig. 1E*). Follow-up tests on the main effect of age indicated that late adolescent females investigated their partner more frequently than early adolescent females [*t* (1,46)=3.52, *p*=0.001] and adult females [t(1.46)=2.23, *p*=0.031]. In contrast to males, mPAE females did not exhibit mPAE-induced changes in social motivation (*Fig. 1F*), social contact (*Fig 1G*), or social play (*Fig. 1H*). In females, the ANOVA revealed a main effect of age on social contact [*F*(2,46)=6.49, *p*=0.0033](*Fig. 1G*), with late adolescent females engaging in more contact with their partners compared to early adolescent [*t*(1,46)=2.83, *p*=0.021] and adult [*t*(2, 46)=3.32, *p*=0.005] females. Female social play also differed as a function of age [*F*(2,46)=9.97, *p*=0.0003],with late adolescents exhibiting more play behavior than early adolescents [*t*(1,46)=2.51, *p*=0.047] and adults [*t*(1,46)=2.51, *p*=0.0002].

Overall, these results demonstrate that mPAE on G12 produced long-lasting social impairments, and that males displayed more mPAE-induced reductions than females, indicating increased overall impairment in males.

### G12 mPAE attenuates the role of the BLA→PL projection in modulating social behavior in males but not females

We next assessed the potential role of the BLA→PL projection in the social impairments produced by G12 mPAE in adult offspring using a viral chemogenetic approach (*Supplementary Figure 1A*). We confirmed that BLA pyramidal cells containing the inhibitory DREADD virus (pAAV-CaMKIIa-hM4D(Gi)-mCherry) were able to be inhibited by CNO using whole-cell patch-clamp electrophysiology (*Supplementary Figure 1B-F*). We then examined the effect of BLA→PL inhibition on adult social behavior by injecting either ACSF (control) or CNO into the PL just before testing across two test days. Interestingly, while we did not find a significant main effect of G12 exposure on social investigation, we did find a main effect of drug injection [*F*(1,14)=15.97, *p*=0.001] and a G12 exposure x drug interaction [*F*(1,14)=5.12, *p*=0.04]. Post-hoc analysis showed that, in air-exposed males, CNO significantly reduced social investigation compared to ACSF [*t*(14)=3.96, *p*=0.003], whereas in mPAE-exposed males CNO had no effect [*t*(14)=1.42, *p*=0.358](*Fig. 2A*). For male social preference, we found no main effect of exposure nor an exposure x drug interaction, but we did find a significant main effect of drug [*F*(1,14)=12.90, *p*=0.003](*Fig. 2B*), with CNO reducing social preference regardless of G12 exposure. There were no other effects of G12 exposure or drug on male contact behavior (*Fig. 2C*) or social play (*Fig. 2D*).

**Figure 2.**
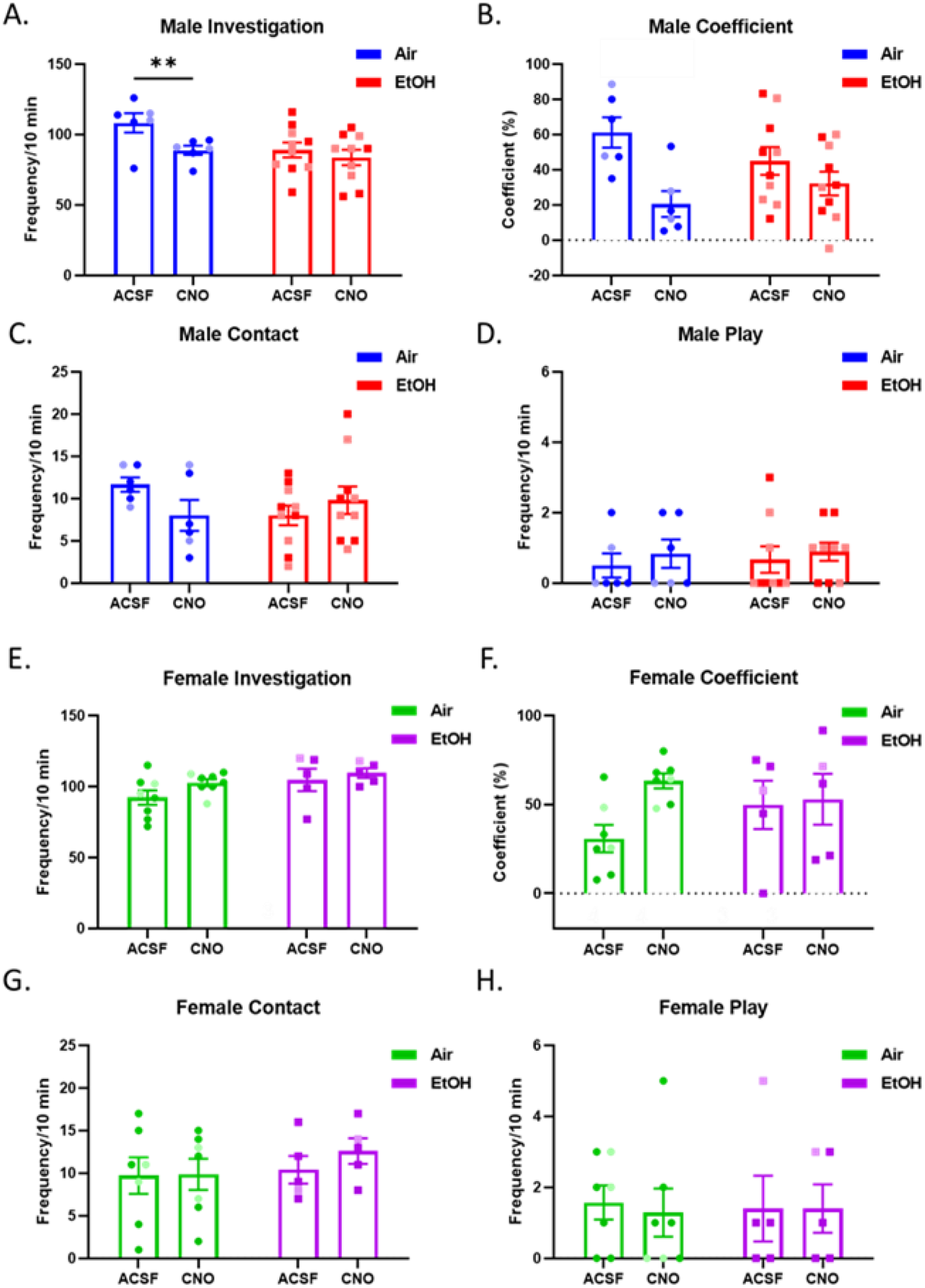
The impact of chemogenetic inhibition of BLA→PL and social behaviors of adult (~P115) male and female rats exposed to air or mPAE) on G12. Air males injected with CNO had reduced **(A)** social investigation compared to air males injected with ACSF, while mPAE males showed no change. Males injected with CNO had reduced **(B)** social preference compared to ACSF-injected males, regardless of exposure. There were no effects in males on **(C)** social contact or **(D)** social play. Females showed no exposure or drug effects on any variable, including **(E)** social investigation, **(F)** coefficient, **(G)** contact, or **(H)** play. ** indicates a significant effect of drug within an exposure group (p−0.05).

Assessment of BLA→PL inhibition on female social behavior revealed no effects of exposure, drug, or interactions on social investigation (*Fig. 2E*), social preference (*Fig. 2F*), contact behavior (*Fig. 2G*), or play (*Fig. 2H*).

Taken together, these results show that chemogenetic inhibition of the BLA→PL modulates social behavior in control males, but its role specifically in social investigation is attenuated by mPAE. Interestingly, female social behavior was not affected by BLA→PL inhibition regardless of prenatal exposure, indicating that the recruitment of the BLA→PL in social behavior may be sex specific.

### CNO injection does not affect social behavior in rats infused with control virus

To confirm that CNO injected into the PL did not have any off-target effects on social behavior, we did a separate analysis comparing ACSF and CNO in air- and mPAE-exposed offspring infused with control virus [(pAAV-CaMKIIa-mCherry). We found no effect of CNO in either males (*Supplementary Fig. 2*) or females (*Supplementary Fig. 3*) indicating that CNO did not impact social behavior in offspring infused with control virus.

### G12 mPAE does not alter glutamate transmission in the PL PL2/3 pyramidal cells

We first examined how mPAE impacted various neurophysiological aspects of PL2/3 pyramidal cells. When assessing the membrane properties of PL2/3 pyramidal cells, we did not find any effects of mPAE on the membrane capacitance or membrane resistance in males or females (*Supplementary Table 1*). When examining glutamate transmission in males, we found no effect of exposure on either sEPSCs (frequency: *Fig. 3B;* amplitude: *Fig. 3C*) or mEPSCs (frequency, *Fig. 3C;* amplitude: *Fig. 3D*). Similarly, in females we found no differences in sEPSCs (frequency: *Fig. 3A;* amplitude: *Fig. 3B*) or mEPSCs (frequency: *Fig. 3C;* amplitude: *Fig. 3D*). Together, these results indicate that neither AP-dependent nor-independent glutamate transmission onto PL2/3 pyramidal neurons was affected by mPAE.

**Figure 3.**
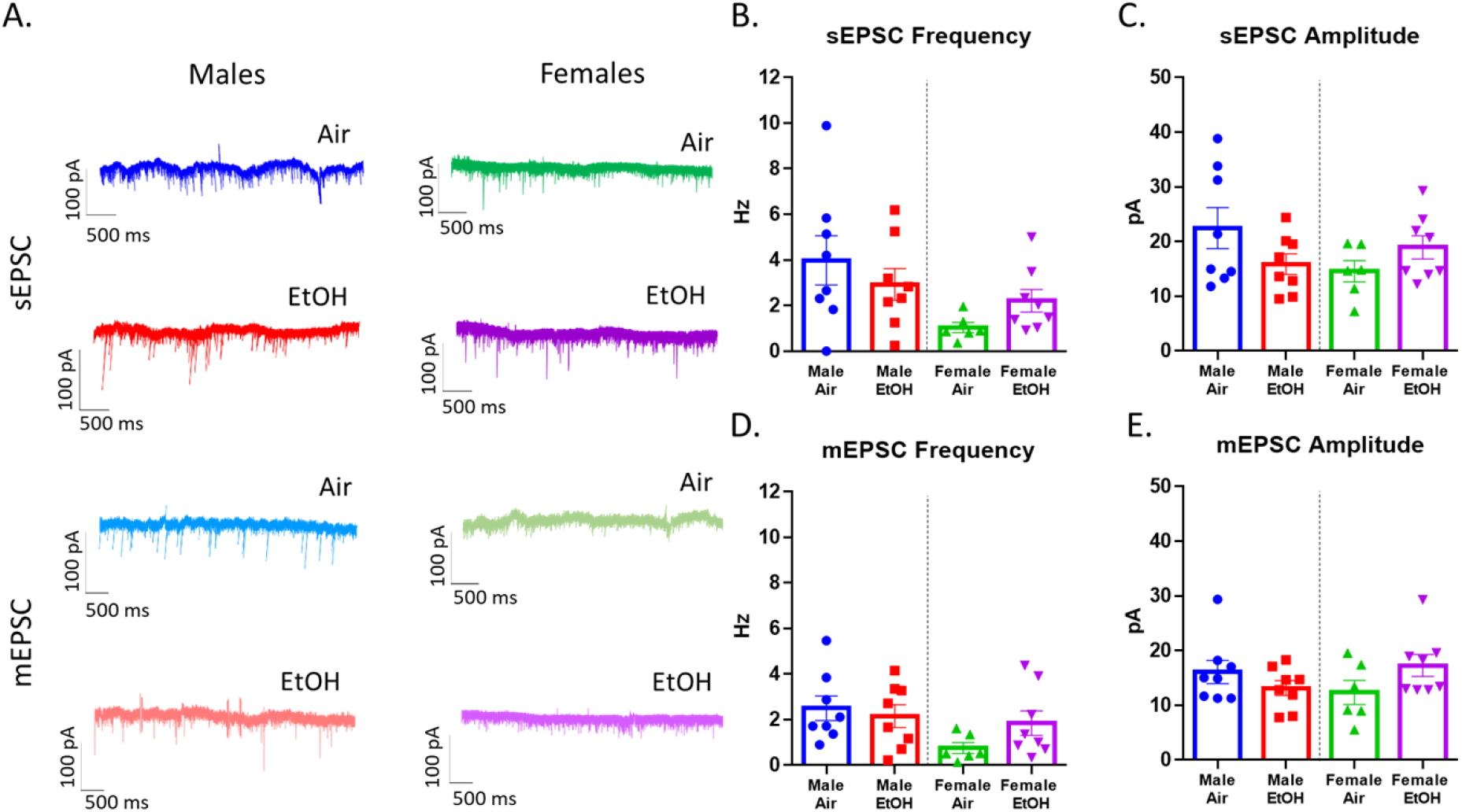
Basal sEPSCs and mEPSCs in PL layer 2/3 pyramidal cells in males and females exposed to air or mPAE on G12. **(A)** Representative sEPSC (top) and mEPSC (bottom) traces from air- or mPAE-exposed adult males (left) and females (right). There were no exposure effects in males or females on sEPSC **(B)** frequency or **(C)** amplitude, nor were there exposure effects on mEPSC **(D)** frequency or **(E)** amplitude of either sex.

### G12 mPAE decreases the excitability of PL pyramidal cells

We next assessed the intrinsic excitability of PL2/3 pyramidal cells by recording RMP followed by AP firing. We found no difference in RMP in males or females (*Supplementary Table 2*). In males, we also found no difference in the rheobase, but there was a significant increase in the rheobase of mPAE females [*t*(10)=3.21, *p*=0.009](*Supplementary Table 1*), indicating that more current was required to fire an AP in mPAE females than control females. When examining the firing patterns of these cells while injecting a series of depolarizing currents, in males we found a significant effect of exposure [*F*(1,170)=4.69, *p*=0.044] and a significant step x exposure interaction [*F*(14, 2265)=2.32, *p*=0.005](*Fig. 4A&C*). When examining the cell properties of the first AP fired, we found an increase in the time to first AP in mPAE males compared to air males, but no effects on any other variable (*Supplementary Table 2*). In females, we also found a significant main effect of exposure [*F*(1,100=6.78, *p*=0.026] and a step x exposure interaction [*F*(14,140)=3.09, *p*=0.0003](*Fig. 4B&D*) on AP firing. However, there were no effects of exposure on any other AP firing variables in females (*Supplementary Table 2*). These firing results indicate that, while all cells increased the number of APs fired per depolarization step, cells from mPAE offspring fired fewer at each step than their air-exposed counterparts. Together these data demonstrate that mPAE decreased the excitability of PL2/3 pyramidal cells, but through different mechanisms in males and females.

**Figure 4.**
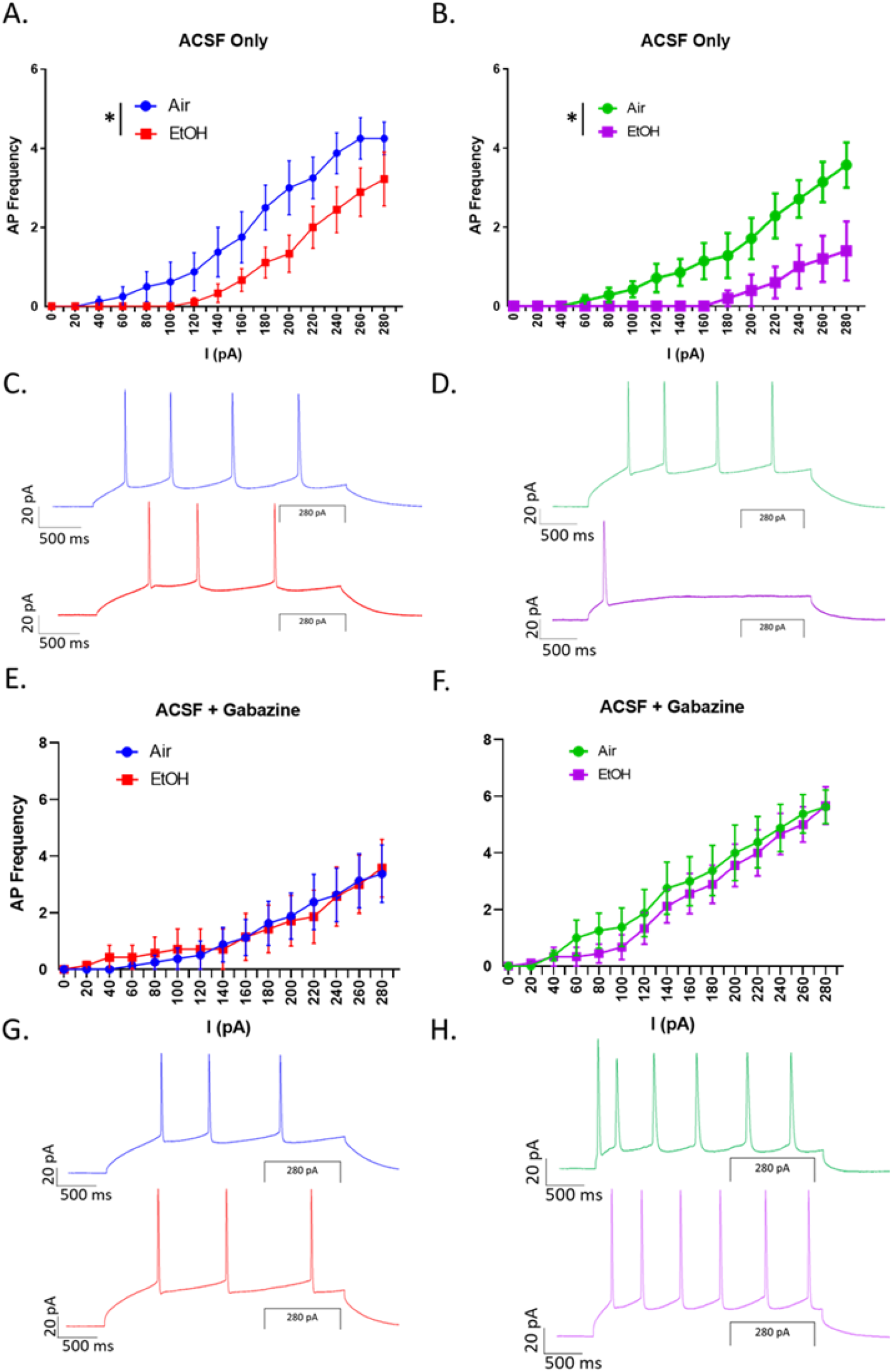
AP firing of PL layer 2/3 pyramidal cells across exposure in males and females exposed to air or mPAE on G12. Increasing current steps (pA) elicited fewer APs in mPAE cells than air cells in **(A)** males and **(B)** females under basal conditions (ACSF only). Representative AP firing traces from **(C)** males and **(D)** females. In the presence of GABA_A_ receptor antagonist gabazine (10 μM), increasing current steps did not elicit a difference in the number of APs fired in either **(E)** males or **(F)** females. Representative traves of AP firing in the presence of gabazine in **(G)** males and **(H)** females. * indicates a significant effect of exposure on the number of APs fired (*p*<0.05)

### Inhibitory modulation of PL pyramidal cell excitability is disrupted following G12 mPAE

Since mPAE effects on PL2/3 cell excitability were not due to alterations in glutamate transmission (*Fig. 3*), to determine whether mPAE altered inhibitory modulation of PL2/3 pyramidal cell excitability, we repeated the AP firing experiments described above in a separate set of neurons in the presence of the GABA_A_ receptor antagonist, gabazine, to block GABAergic inhibition of PL2/3 cells. In contrast to what we observed with basal firing, in the presence of gabazine we observed no effect of exposure or no current injection x exposure interaction in males (*Fig. 4E*) or females (*Fig. 4F*). Assessment of the first AP also showed no effects of exposure in males (*Supplementary Table 3*). In females, we found only that there was a significant increase in the peak of the first AP in mPAE cells compared to air cells (*Supplementary Table 3*). Together, these results indicate that gabazine application eliminated preexisting differences in AP firing between cells from air- and mPAE-exposed offspring, suggesting mPAE may increase inhibitory modulation of AP firing in the PL.

## Discussion

The goals of this investigation study were to determine whether G12 mPAE could produce long-term social impairments like binge-level PAE, and to examine neuroadaptations underlying the observed social impairments in adults. We discovered that G12 mPAE produced social impairments that lasted into adulthood, with males exhibiting impairments on the majority of social behaviors examined, while females showed impairment only on social investigation. We then tested the hypothesis that mPAE increases BLA→PL activity, contributing to these social impairments, by chemogenetically inhibiting this projection during social testing. We found that BLA→PL inhibition reduced social behavior in control males while having no effect in mPAE males or females of either exposure group. Finally, given the role of the PL in social suppression and avoidance, we examined mPAE-induced neurophysiological changes in PL2/3 pyramidal cells. We found that mPAE decreased the intrinsic excitability of these cells in both sexes, potentially due to increased inhibitory suppression of pyramidal cell excitability.

G12 is a well-established vulnerable timepoint for PAE, with several studies showing behavioral impairments resulting from G12 PAE. More specifically, extreme binge-level PAE (BECs ranging from 280-400 mg/dL) on G12 led to reductions of social investigation, social preference, and social contact through adulthood in male offspring [11, 14, 62], the same social behaviors impacted in the present work. We also found that females were less affected by G12 mPAE, exhibiting fewer social impairments than males, consistent with what has been reported following extreme binge-level PAE [11, 13, 14, 16, 62, 63]. Fewer studies have examined the impact of mPAE on social behavior, but those studies have shown that regardless of the timing of exposure, male social behavior is often more affected than female social behavior [25, 64]. We have also shown that G12 mPAE enhances non-social anxiety-like behavior in adolescent males, but not adolescent females [32]. Thus, our current findings align with previous work showing that mPAE significantly impairs social behaviors long-term, with males appearing more vulnerable than females.

Previous work showed that PAE increases BLA activity [48, 49] and that activation of the BLA→PL leads to social avoidance and other fear/anxiety-related behaviors [47, 62, 65, 66]. Thus, we chemogenetically inhibited the BLA→PL in adult offspring during social testing with the goal of reversing social deficits in mPAE offspring. In contrast to our expectation, we found that air-exposed males exhibited reduced social investigation following BLA→PL inhibition, while neither mPAE males nor females of either exposure group were affected by BLA→PL inhibition. The lack of BLA→PL inhibition effects on social behavior in mPAE males may indicate that mPAE blunts the role of the BLA→PL in modulating male social suppression and social avoidance behavior. This is consistent with work showing that early-life stress weakened BLA÷mPFC inputs and reduced their influence on behavior [67], suggesting that developmental insults may generally attenuate BLA→mPFC modulation of behavior. However, there are other possibilities that should be considered. BLA projections to the PL have numerous targets, including different neuron types within different PL layers that each have distinct behavioral implications [65, 68–71]. Our chemogenetic approach targeted all BLA terminals within the PL, and whether and how these different projections interact within the mPFC microcircuitry to regulate behavior, and are influenced by PAE, is unknown. Therefore, more work is necessary to fully understand the role of these circuits in modulating normal and pathological social behaviors. Despite this, our findings provide a novel assessment of how mPAE may disrupt normal BLA→PL modulation of social behavior.

A noteworthy observation is that inhibiting the BLA→PL in control males reduced social investigation, contrasting with what has been previously shown in mice [47]. Methodological differences between these two studies, such as the age and species of laboratory animal and the specific approach used to manipulate the circuit (optogenetic vs. chemogenetic) may explain the opposing results. Overall, our finding that this manipulation reduced social investigation in control males adds complexity to our ongoing exploration of the neural circuitry underlying social behavior and confirms that different factors such as age, species, and developmental experiences are important variables to consider when examining these circuits.

Interestingly, female social behavior was not affected by BLA→PL inhibition, regardless of prenatal exposure condition. While no studies have examined this specific pathway during female social behavior, previous work has found that this projection is less active in females and has less control over PL activity than in males, potentially due to circulating sex hormones [72]. Together with the present work, this suggests that females may recruit the BLA→PL to a lesser degree than males for behavioral control. This may be particularly relevant for social interaction, with other work showing less activation of the BLA and PL during social learning [43] and functional connectivity between the amygdala and PFC in a social context [73] in females than in males. Thus, our findings support the idea that the BLA→PL may be less important for social interaction in females, which could be one mechanism by which females are less vulnerable to the socially impairing effects of PAE.

Finally, we assessed whether G12 mPAE would functionally alter PL2/3 pyramidal cells. Previous studies found that PAE alters mPFC neuron morphology, numbers, and function, and mPFC network connectivity [23, 24, 50–53, 74–76]. Given the role of the PL in modulating negative social behaviors, we aimed to determine how G12 mPAE affected PL physiology, as this has not been previously examined. We found that mPAE reduced the excitability of PL2/3 pyramidal cells, which has been observed across various PAE exposure paradigms, test ages, and cell types in other mPFC layers. For example, PAE increased inhibitory input onto mPFC layer 5 and layer 6 pyramidal cells which was related to reduced excitability of those cells [50, 52, 75]. Our results are consistent with this work and extend it to layer 2/3, an important contribution to a growing interest in how developmental insults lead to cortical dysfunction. Furthermore, consistent with a PAE-induced increase in inhibitory interneuron populations and GABAergic inhibition [50, 53, 75], mPAE-induced differences in pyramidal cell excitability were abolished when blocking inhibitory transmission in the present work, suggesting that the inhibitory drive onto these pyramidal cells may be potentiated following mPAE. A direct examination of changes to mPFC interneurons, GABAergic transmission, and impacts to the larger mPFC microcircuitry and related behaviors is required to fully support this conclusion.

Taken together, this study provided valuable insight into the long-lasting neuroadaptations caused by G12 mPAE, particularly in specific brain regions/circuits that are implicated in social deficits like those associated with FASD. We observed several sex-specific including baseline differences in mPAE-induced social impairments and the role of the BLA→PL in modulating social behavior, which provides a framework for further mechanistic investigations focused on sex as a contributing variable. Despite its limitations, this work also set a foundation for several future studies that will enhance our understanding of the specific neuronal cell types and circuits that are affected by PAE, paving the way to examinations of specific manipulations and treatments to improve health outcomes in individuals with FASD across the lifespan.

## Acknowledgements

The authors would like to acknowledge and thank Drs. Molly Deak, David Werner, and Kathryn Lanza for their instruction on numerous technical aspects of this work, and undergraduate assistants Michelle Montero, Clare Wiberg, and Dana Silberstein for their contributions to data analysis and tissue preparation. The authors have no conflicts of interest to disclose. This work was supported by NIAAA grants P50 AA017823 and R21 AA027873, as well as grants to the author from the Research Society on Alcoholism and the American Psychological Association.

## Author Contributions

KRP: Conceptualization, methodology, formal analysis, investigation, writing-original draft, writing- reviewing & editing, visualization, and funding acquisition. MBS: investigation, writing-reviewing & editing. JMJ: investigation. EIV: conceptualization, methodology, formal analysis, investigation, writing-reviewing & editing, supervision. MRD: conceptualization, methodology, writing- reviewing & editing, supervision, project administration, funding acquisition

## Supplementary Materials

### Supplementary Methods

#### Subjects

Male and female Sprague-Dawley rats bred in our colony were used for all experimental procedures. Upon weaning, animals were pair-housed with ad libitum access to food (Picolab Laboratory Rodent Diet 5L0D, LabDiet, St. Louis, MO) and water. Rats were maintained on a 12-hour light/dark cycle beginning at 0700 in a temperature-controlled vivarium. Early adolescent (postnatal day (P) 28), late adolescent (P42), and adult (P77) rats were used for baseline social behavior testing as described below, and only adult (P70+) rats were used for chemogenetic manipulation experiments and electrophysiology. All procedures were conducted in accordance with protocols that were approved by the Binghamton University Institutional Animal Care and Use Committee.

#### Breeding and Moderate Prenatal Alcohol Exposure

For breeding, two adult female breeders were placed in a large cage with one adult male breeder (Envigo, Indianapolis, IN). Each morning for four days, the females were checked via vaginal smear for the presence of sperm. The day of sperm detection was designated as G1. Upon confirmation of pregnancy, females were single housed for the duration of pregnancy. Pregnant dams received high-nutrient food (Purina 5008C33 Rat Chow, LabDiet, St. Louis, MO) and water ad libitum, and were weighed on G1, G10, and G20 to monitor weight-gain. On G12, dams were transferred to a clean cage, which was placed into one of two vapor inhalation chambers as previously described (Rouzer et al., 2017). From 0900 to 1500, dams were exposed to either room air or vaporized ethanol. This exposure paradigm has been demonstrated to produce a peak blood ethanol concentration in the moderate range (60-90 mg/dl (Rouzer et al., 2017)). To minimize exposure to ethanol that may have been absorbed into the food or bedding, these were replaced upon completion of the vapor exposure, and dams were returned to the colony room. After giving birth, pups remained with the dam until the day of weaning (P21) and pup weights were obtained on P2, P7, and P12.

#### Social Interaction Testing

Social behavior was assessed in early adolescent, late adolescent, and adult male and female rats that had undergone our G12 exposure paradigm, and each rat was only tested for social behavior once. All social behavior testing was performed between 0900 and 1100 and was conducted as previously described (Diaz et al., 2016). The testing chambers (30 x 20 x 20 cm for adolescents; 45 x 30 x 30 cm for adults) contained fresh woodchip bedding, were composed of Plexiglas and were divided into two equal-sized compartments by a clear Plexiglas partition with an aperture (7 x 5 cm for late adolescents and 9 x 7 cm for adults) to allow test rats to move freely between the two compartments.

On the day of testing, experimental rats were removed from their home cages, weighed, and placed into the testing chamber for a habituation period of 30 minutes. Following the habituation period, an experimentally naïve partner was placed into the chamber for a ten-minute test period. Social partners were all unfamiliar to the experimental rat and the experimental rat was always heavier than the partner. Following testing, the experimental rats and partners were returned to their home cages.

All behaviors were video recorded and saved for later scoring of social interaction by a researcher blind to the exposure condition. Behaviors scored included frequencies of social investigation (sniffing of the partner), contact behavior (grooming of and climbing over/under the partner), and play behavior (pouncing, following/chasing, pinning). Social motivation via social preference/avoidance was measured as crossovers from one compartment of the testing apparatus toward (preference) or away from (avoidance) the partner, and a social preference/avoidance coefficient was calculated [coefficient (%) = (crossovers toward partner – crossovers away from the partner)/(total crossovers toward/away) x 100].

#### Data Analysis

Behavioral data were analyzed using Prism 6 (GraphPad). Previous literature using the modified social interaction test has indicated that there are overall differences in the social-behavioral profiles of males and females (Vetter-O’Hagen & Spear, 2012; Varlinskaya, Truxell, & Spear, 2014; Kim & Spear, 2016); therefore, data from male and female animals were separated throughout social behavioral assessment. Statistical analyses were conducted as two-way between subjects ANOVAs comparing across age and exposure group. Following a significant main effect of age or significant exposure x age interaction, Bonferroni corrected post hoc tests were conducted to detect significant differences between groups. The significance threshold was set at p < 0.05, and data are represented as mean ± SEM.

#### Viral Infusion and Cannulation Surgery

Adult (P70-80) offspring were anesthetized with isoflurane (2-4%) and placed in a stereotactic apparatus (David Kopf Instruments, Tujunga, CA). A midline incision and craniotomies were made for viral transduction into the BLA. For viral infusions into the BLA, the coordinates used were AP: +2.5, ML: ±5.0, and DV: −8.8 mm relative to Bregma (Paxinos & Watson, 2014). A 10 μL Hamilton syringe with a 26-gauge needle was used to deliver viral solutions into the BLA at a rate of 0.1 μL/min using a microsyringe pump (Stoelting). An additional 5 minutes was allowed for the virus to diffuse into the tissue, at which point the injection needle was slowly removed. Rats were randomly assigned to receive either an inhibitory chemogenetic virus (pAAV-CaMKIIa-hM4D(Gi)-mCherry; Addgene, Watertown, MA) or a control virus containing only the attached fluorophore (pAAV-CaMKIIa-mCherry).

Following viral infusion, an additional craniotomy was made for implantation of a bilateral guide cannula (26 gauge; Plastics One, Roanoke, VA) into the PL. Cannulae terminated 1 mm above the PL using the following coordinates: AP: +3.5, and DV: −2.5 relative to Bregma. Because of the PL’s proximity to the midline, a single bilateral cannula with a distance of 1 mm between guides was implanted on the midline. Dental cement (Lang Dental, Wheeling, IL) was affixed to the guide cannula and 3-4 bone screws were used to anchor and stabilize the cement cap. To maintain patency of the guide during recovery, a 10.5 mm cannula dummy was inserted into the guide, and a dust cap was placed over the top of the dummy cannula. At the end of surgery, rats were treated with topical antibacterial cream to prevent infection at the incision site, and they received injections (i.p.) of buprenorphine (0.03 mg/kg) for pain management every 12 hours for 48 hours after surgery. Post-operative care was conducted once per day for 10 days after surgery, and twice per week thereafter until testing.

#### Microinjection Procedure and Social Interaction Testing

Social behavior testing began at least six weeks after viral infusion surgery to allow viral transduction from the site of infusion at the BLA to BLA terminals within the PL, and behavior testing followed the same procedures as described above for adult rats. Five days prior to the first social interaction testing day, rats were extensively handled and gradually habituated to the manipulations associated with the microinjection procedure. On the day of testing, rats received a microinjection into the PL of either artificial cerebrospinal fluid (ACSF) or 300 μM clozapine N-oxide (CNO; Hello Bio). For the microinjections, dummy cannulas were removed and a 33-gauge bilateral internal injector cannula was inserted into the guide cannula such that the injector extended 1 mm below the end of the guide. Drug solutions were infused into the PL over a 1-minute period with a total of 0.5 μL volume delivered per side, and injectors were left in place for one additional minute to allow drug diffusion into the tissue.

Following the injection procedure, rats were immediately placed into the social interaction chambers and testing began as described above. To determine whether the CNO injection influenced locomotor behavior in a nonsocial and a social setting, we also included these measures as total crosses between compartments during the habituation and interaction phases, respectively, in our assessment.

#### Experimental Design

Social behavioral testing with chemogenetic manipulation was designed as a 2 (exposure) x 2 (virus) x 2 (drug) mixed experimental design, with exposure and virus groups as between subjects variables. The drug microinjection groups were a within subjects variable, such that each rat was tested twice, once receiving ACSF and once receiving CNO. The two test days occurred 48 hours apart with a novel social partner each time, and the order of drug delivery was randomized to eliminate order effects of multiple test procedures. A schematic outlining this experimental design can be found in Supplementary Figure 1A.

#### Verification of Virus/Cannula Placement

Following social testing, rats were sacrificed using a 180 mg/kg dose of sodium pentobarbital and transcardially perfused with cold 4% paraformaldehyde. Brains were extracted, fixed in 4% paraformaldehyde for 12-16 hours, and bathed in 30% sucrose. Coronal slices (40 μm) were made using a cryostat (CM1510, Leica Biosystems) to confirm placement of guide cannulae over the PL and viral injections into the BLA. Slides containing the PL were thaw-mounted onto charged slides and placement of guide cannulae was confirmed using a standard optical microscope. Any data obtained from animals with a misplaced guide cannula was retained for potential control studies but removed from data analysis. Slices containing BLA were preserved in antifreeze for immunofluorescence procedures.

To verify virus placement into the BLA, free floating BLA-containing slices were washed three times for 10 min each in phosphate buffered saline (PBS), followed by a 1 hr. incubation in blocking buffer containing 10% Normal Donkey Serum, 1% Bovine Serum Albumin, glycine (300 mM), 10X PBS, and 0.2% Triton X-100. Sections were then incubated overnight at 4°C in primary antibody: mAb mouse anti-mCherry (632543, Takara Bio, San Jose, CA, 1:1000 dilution). After washing 3 × 10 min in PBS, sections were incubated for 2 hr. at room temperature in secondary antibody: pAb donkey anti-mouse: Alexafluor 594 (715-585-151, Jackson Immunoresearch Laboratories Inc., West Grove, PA, USA, 1:2000 dilution). Sections were mounted using Prolong Gold Antifade Mountant with DAPI (Thermo Fisher Scientific, Waltham, MA) and allowed to air dry for at least 24 hr. Fluorescent images were collected on a BX-X800 fluorescent microscope (Keyence, Osaka, Japan) at 2X magnification. A field-of-view image was taken through DAPI and Texas Red filters to visualize both the BLA and the location of the virus. Data were analyzed from animals in which there was a unilateral or bilateral BLA hit, with a hit defined as at least half the area of the BLA showing presence of the virus.

#### Data Analysis

For analysis of behavioral data in offspring who received the active inhibitory DREADD virus. two-way mixed ANOVAs were conducted to compare across exposure and drug injection, with exposure as a between subjects variable and drug injection as a repeated measures variable. A subset of rats received an infusion of control virus to confirm that any observed CNO effects were due to the presence of the inhibitory DREADD virus, so a separate paired t-test analysis was conducted within each exposure group to confirm that CNO had no effect in these animals. Following a significant exposure x drug interaction, Bonferroni corrected post hoc tests were conducted to detect significant differences between groups. The significance threshold was set at p < 0.05, and data are represented as mean ± SEM.

#### Slice Preparation

PL-containing brain slices were prepared as described previously (Przybysz, Werner, & Diaz, 2017). Rats were deeply sedated with isoflurane (4-5%) and quickly decapitated. Brains were rapidly removed and submerged in cold oxygenated (95% O_2_–5% CO_2_) N-methyl-d-glucamine (NMDG) cutting solution containing (in mM): NMDG (135), KCl (1), KH_2_PO_4_ (1.2), MgCl_2_*6H_2_O (1.5), CaCl_2_*2H_2_O (0.5), 80% choline bicarbonate (35.4), dextrose (10), and ketamine (0.43). PL-containing coronal slices (300-350 μm) were made using a Vibratome (Leica Microsystems). Slices were incubated at 34°C for a recovery period of at least 40 minutes in normal ACSF containing (in mM): NaCl (126), KCl (2), NaH_2_PO_4_ (1.25), NaHCO_3_ (26), glucose (10), CaCl_2_ (2), MgSO_4_ (1), ascorbic acid (0.4), continuously bubbled in 95% O_2_–5% CO_2_. All experiments were conducted approximately 1-4 hours after slice preparation.

#### General Electrophysiology

For whole-cell patch-clamp recordings, neurons were visualized using infrared interference-contrast microscopy. Data were acquired with a MultiClamp 700B (Molecular Devices, Sunnyvale, CA) at 10 Hz, filtered at 1 Hz, and stored for later analysis using pClamp software (Molecular Devices). Cells were visualized using infrared-differential interference microscopy (Olympus America, Center Valley, PA). Recordings were made from pyramidal cells in Layer 2/3 of the PL, initially identified based on morphology and membrane properties (pyramidal cells: triangle-shaped with a visible apical dendrite, membrane capacitance > 150 pF, membrane resistance < 150 MΩ). Cell type was confirmed *post hoc* by examining the resting membrane potential and pattern of action potential firing (Ji, 2012; Povysheva et al., 2006; van Aerde & Feldmeyer, 2015). Recordings were collected with patch pipettes filled with a K-gluconate-based internal solution containing (in mM): K-gluconate (120), KCl (15), EGTA (0.1), HEPES (10), MgCl_2_ (4), MgATP (4), Na_3_GTP (0.3), Phosphocreatine (7), and QX-314 Br (1.5), pH: 7.3; mOsm: 300. Access resistance was monitored throughout each experiment, and any cells in which the access resistance changed by > 20% were removed from analysis. Three types of experiments were conducted, as described below.

##### Basal Intrinsic Excitability

Intrinsic excitability of pyramidal cells was assessed in a current-clamp configuration at resting membrane potential (RMP) in a bath containing only ACSF. Cells were opened in voltage-clamp configuration and immediately switched to zero-current configuration to obtain a reading of the RMP of the cell. Cells were then switched to current-clamp configuration and allowed to dialyze for at least 5 minutes. A series of depolarizing current steps (increasing steps of 10 pA beginning at −100 pA, 500 ms step duration) was applied and the following parameters were assessed: the amount of current required to fire an action potential (rheobase), the membrane potential at which the first action potential fired, the time to first action potential, and the number of action potentials fired with increasing current steps.

##### GABAergic Modulation of Firing

To assess the role of GABAergic interneurons in modulating pyramidal cell firing, identical experiments were conducted as above in a separate subset of neurons. Following the basal ACSF recording, a second identical recording was conducted while pharmacologically blocking GABAA receptors with gabazine (10 μM).

##### Spontaneous Glutamate Transmission

Following the assessment of intrinsic excitability, spontaneous excitatory postsynaptic currents (sEPSCs) were recorded in voltage-clamp configuration. Baseline glutamate transmission was recorded during the continued application of gabazine (10 μM). Miniature (m) EPSCs were then recorded in the presence of tetrodotoxin (0.5 μM) to assess action potential-independent transmission.

#### Drugs and Chemicals

Following the assessment of intrinsic excitability, spontaneous excitatory postsynaptic currents (sEPSCs) were recorded in voltage-clamp configuration. Baseline glutamate transmission was recorded during the continued application of gabazine (10 μM). Miniature (m) EPSCs were then recorded in the presence of tetrodotoxin (0.5 μM) to assess action potential-independent transmission.

#### Data Analysis

Intrinsic excitability recordings were analyzed using Clampfit (pClamp), and EPSC recordings were analyzed using MiniAnalysis (SynaptoSoft Inc.). All data were statistically analyzed with Prism 6, and males and females were analyzed separately. Statistical analyses for excitability variables of rheobase, membrane potential at the first action potential, and time to first action potential were all conducted as t-tests between exposure groups. The number of action potentials fired with increased depolarization steps was initially analyzed as a mixed twoway ANOVA across depolarization step (repeated measures) and exposure group, with a significant main effect of exposure and/or a significant interaction indicating differences in excitability between exposure groups. Gabazine recordings were analyzed similarly, and comparisons between ACSF and gabazine firing were also conducted as two-way ANOVAs (repeated measures for both depolarization step and drug applied) within each exposure group. sEPSC and mEPSC recordings were also analyzed as t-tests between exposure groups. As with behavioral experiments, the significance threshold was set at p < 0.05. Data are represented as mean ± SEM.

### Supplementary Results

#### G12 mPAE produces sex-specific social impairments lasting into adulthood

When examining the variable of social investigation, we found a significant main effect of G12 exposure in males [*F*(1,46) = 10.12, *p* = 0.0026], but no main effect of age [*F*(2, 46) = 1.395, *p* = 0.092], or an age x exposure interaction [*F*(2, 46) = 0.07, *p* = 0.932] (*Fig. 1A*). Overall, these data indicate that offspring exposed to mPAE on G12 exhibited reduced social investigation, regardless of age. We also examined the social preference/avoidance coefficient in males, which is considered a proxy for social anxiety-like behavior. Our analysis revealed a significant main effect of exposure on this variable [*F*(1,46) = 15.16, *p* = 0.0003] (*Fig. 1B*) with mPAE males exhibiting reduced social motivation compared to air-exposed males. There was no main effect of age [*F*(2,46) = 0.199, *p* = 0.836] nor was there an age x exposure interaction [*F*(2,46) = 0.18, *p* = 0.836]. Social contact behavior, which includes grooming of the partner and climbing over or under the partner by the experimental rat, was also assessed. In males, we found a significant main effect of exposure [*F*(1,49) = 7.68, *p* = 0.0079], with no main effect of age [*F*(2, 49) = 3.04, *p* = 0.057], and no age x exposure interaction [*F*(2, 49) = 0.33, *p* = 0.720) (*Fig. 1C*). Like our other findings in males, mPAE males exhibited reduced contact with their partners regardless of age. Unlike the other social behavior variables, for social play in males we found a found a significant main effect of age [*F*(2, 46) = 6.79, *p* = 0.003]. Follow-up analyses indicated that social play decreased across ontogeny, with adults exhibiting less play than early adolescents [*t*(1, 46) = 3.56, *p* = 0.0056] and late adolescents [*t*(1, 46) = 2.55, *p* = 0.0422] (*Fig. 1D*). There was no main effect of exposure for social play [*F* (1, 46) = 2.14, *p* = 0.15], nor was there an age x exposure interaction [*F*(2, 46) = 0.19, *p* = 0.82].

When examining female social behavior, we found a significant main effect of G12 exposure on social investigation [*F*(1,46) = 19.28, *p* < 0.0001], as well as a significant main effect of age [*F*(2, 46) = 6.376, *p* = 0.0036], but no age x exposure interaction [*F*(2, 46) = 0.092, *p* = 0.91] (*Fig. 1E*). These data indicate that mPAE reduced social investigation in females. Follow-up tests on the main effect of age indicated that late adolescent females engaged in social investigation with their partner more frequently than early adolescent females [*t* (1, 46) = 3.52, *p* = 0.001] and adult females [t(1.46) = 2.23, *p* = 0.031]. When assessing the social preference/avoidance coefficient in females, we found no main effect of exposure [*F*(1,46) = 0.65, *p* = 0.425], no main effect of age [*F*(2,46) = 1.26, *p* = 0.292], and no age x exposure interaction [*F*(2,46) = 0.64, *p* = 0.531] (*Fig. 1F*). In contrast to our findings in males, this indicates that female social preference was not influenced by age or mPAE exposure. Analysis of social contact behavior in females revealed no main effect of exposure [*F*(1, 46) = 0.24, *p* = 0.625] or an age x exposure interaction [*F*(2,46) = 0.28, *p* = 0.754]; we did find a significant main effect of age [*F*(2, 46) = 6.49, *p* = 0.0033] (*Fig. 1G*), indicating that females exhibited agespecific changes to social contact. Follow-up analyses indicated that late adolescent females displayed increased contact with their partners compared to early adolescent [*t*(1, 46) = 2.83, p = 0.021] and adult [t(2, 46) = 3.32, p = 0.005] females. When we examined social play in females, we found a significant main effect of age [*F*(2, 46) = 9.97, *p* = 0.0003] (*Fig. 1H*), with follow-up tests demonstrating that late adolescent females displayed more social play than early adolescents [*t*(1,46) = 2.51, *p* = 0.466] and adults [*t*(1,46) = 2.51, *p* = 0.0002]. There was no main effect of exposure [*F*(1,46) = 0.58, *p* = 0.451], nor was there an age x exposure interaction [*F*(2,46) = 0.05, *p* = 0.95] in female social play (*Fig. 4D*), indicating that mPAE exposure did not influence female play behavior.

Overall, our results from Experiment 1 demonstrate that mPAE on G12 produced long-lasting social impairments, and that males displayed more mPAE-induced reductions than females, indicating increased overall impairment in males.

#### G12 mPAE attenuates the role of the BLA→PL projection in modulating male social behavior

Prior to beginning social behavior testing with experimental offspring, we conducted a pilot study to verify that BLA pyramidal cells containing the inhibitory DREADD virus (pAAV-CaMKIIa-hM4D(Gi)-mCherry) were able to be inhibited by CNO. To do this, we used whole-cell patch-clamp electrophysiology to record the intrinsic excitability of fluorescently labeled (virus-containing) BLA pyramidal cells at baseline and following application of CNO (10 μM). There was a significant effect of CNO on rheobase [*t*(3) = 7.79, *p* = 0.004], indicating that CNO increased the amount of stimulation required to fire an action potential (*Supplementary Figure 1B-F*). When examining the number of action potentials elicited by a series of depolarizing current injections, we found a trend toward a significant main effect of drug [*F*(1, 3) = 7.72, *p* = 0.06], indicating that fewer action potentials were fired in the presence of CNO. There was no drug x current injection interaction [*F*(14, 42) = 0.64, *p* = 0.82] (*Fig. 5D*). Together, these pilot data show that CNO does inhibit neurons containing the DREADD virus following a 6-week incubation period.

To examine the role of the BLA→PL in social behavior, we infused either an inhibitory DREADD virus (pAAV-CaMKIIa-hM4D(Gi)-mCherry) or a control virus lacking the inhibitory DREADD receptor (pAAV-CaMKIIa-mCherry) into the BLA. Six weeks later, we injected CNO into the PL (See *Fig. 6* for a summary of cannula placements) to inhibit BLA terminals in the PL, or ACSF control, just before social interaction testing over two test days (repeated measures across drug injection) in air- or mPAE-exposed adult offspring.

##### Males

When analyzing male social investigation, we did not find a significant main effect of exposure [*F*(1, 14) = 2.60, *p* = 0.13]. However, we did find a main effect of drug injection [*F*(1, 14) = 15.97, *p* = 0.001] and an exposure x drug interaction [*F*(1, 14) = 5.12, *p* = 0.04]. Post hoc analysis of these data showed that, in air-exposed males, CNO significantly reduced social investigation compared to ACSF [*t*(14) = 3.96, *p* = 0.003], whereas in mPAE-exposed males CNO had no effect [*t*(14) = 1.42, *p* = 0.358](*Fig. 2A*). For social motivation in males, we also found no main effect of exposure [*F*(1, 14) = 0.07, *p* = 0.798] nor an exposure by drug interaction [*F*(1, 14) = 3.48, *p* = 0.08]. However, we did find a significant main effect of drug [*F*(1, 14) = 12.90, *p* = 0.003], indicating that CNO reduced social motivation regardless of exposure. On the measure of male contact behavior, we found no main effect of exposure [*F*(1, 14) = 0.41, *p* = 0.533], no main effect of drug [*F*(1, 14) = 0.35, *p* = 0.565], and no exposure x drug interaction [*F*(1, 14) = 2.98, *p* = 0.106], together indicating that neither exposure nor CNO affected social contact behavior in males (*Fig. 2C*). We also found no effects on social play, with no main effect of exposure [*F*(1, 14) = 0.08, *p* = 0.784], no main effect of drug [*F* (1, 14) = 0.86, *p* = 0.372], and no exposure x drug interaction [*F*(1, 142) = 0.03, *p* = 0.856] (*Fig. 2D*). Together, these results indicate that inhibiting the BLA→PL projection in adult males may reduce social behavior, and that exposure to G12 mPAE attenuates this function given that there were no effects of CNO in mPAE males.

##### Females

We also assessed the role of BLA→PL inhibition and mPAE on female social behavior, an analysis that to our knowledge has not been previously conducted. When examining social investigation, we found a trend toward a significant main effect of exposure [*F*(1, 11) = 4.41, *p* = 0.067], no main effect of drug [*F*(1, 11) = 2.21, *p* = 0.165] and no exposure x drug interaction [*F*(1, 11) = 0.03, *p* = 0.591](*Fig. 2E*). For the social preference/avoidance coefficient, we found no main effect of exposure [*F*(1, 11) = 0.25, *p* = 0.630], no main effect of drug [*F*(1, 11) = 2.11, *p* = 0.174] and no exposure x drug interaction [*F*(1,11) = 1.21, *p* = 0.294](*Fig. 2F*). On the variable of social contact behavior, we also found no main effect of exposure [*F*(1, 11) = 0.96, *p* = 0.349], no main effect of drug [*F*(1, 11) = 0.749, *p* = 0.405], and no exposure x drug interaction [*F*(1, 11) = 0.749, *p* = 0.405](*Fig. 2G*). There was also no main effect of exposure [*F*(1, 11) = 0.002, *p* = 0.962], no main effect of drug [*F*(1, 11) = 0.95, *p* = 0.567], and no exposure x drug interaction [*F*(1, 11) = 0.95, *p* = 0.567] for female social play.

Taken together, our results from Experiment 1 show that the BLA→PL may play a role in modulating social behavior in control males, but this role is attenuated following mPAE. Interestingly, inhibiting the BLA→PL did not alter female social behavior regardless of prenatal exposure, indicating that the recruitment of the BLA→PL in social behavior may be sex specific.

#### CNO injection does not affect social behavior in rats infused with control virus

To confirm that CNO does not have any off-target effects on social behavior when injected into the PL, we did a separate analysis comparing ACSF and CNO in air- and mPAE-exposed offspring infused with control virus [(pAAV-CaMKIIa-mCherry). In males, there was no effect of CNO on any variable, including investigation [Air: *t*(7) = 1.39, *p* = 0.206; EtOH: *t*(11) = 0.29, *p* = 0.781], coefficient [Air: *t*(7) = 0.79, *p* = 0.455; EtOH: *t*(11) = 1.21, *p* = 0.250], contact [Air: *t*(7) = 0.28, *p* = 0.787; EtOH: *t*(11) = 0.29, *p* = 0.776], or play [Air: *t*(7) = 1.26, *p* = 0.249; EtOH: *t*(11) = 0.96, *p* = 0.358](*Supplementary Fig. 2*). Similarly, CNO also had no impact on social behavior in females injected with control virus. Specifically, there was no effect of CNO on investigation [Air: *t*(5) = 270, *p* = 0.798; EtOH: *t*(6) = 0.65, *p* = 0.541], coefficient [Air: *t*(5) = 0.41, *p* = 0.693; EtOH: *t*(6) = 1.37, *p* = 0.219], contact [Air: *t*(5) = 1.14, *p* = 0.299; EtOH: *t*(6) = 0.82, *p* = 0.442], or play [Air: *t*(5) = 1.51, *p* = 0.182; EtOH: *t*(6) = 0.57, *p* = 0.599](*Supplementary Fig. 3*). Together, these data indicate that CNO did not impact social behavior in offspring infused with control virus.

#### G12 mPAE does not alter the membrane properties of PL layer 2/3 pyramidal cells

Experiment 3 was designed to examine physiological changes to PL function following mPAE. When assessing potential changes to the membrane properties of PL layer 2/3 pyramidal cells in males, we did not find any differences on the membrane capacitance [t(13) = 0.33, p = 0.743] or membrane resistance [t(13) = 0.2, p = 0.804](*Supplementary Table 1*). We also did not find changes to these variables in females [capacitance: t(10) = 0.78), p = 0.455; resistance: t(10) = 1.57, p = 0.147](*Supplementary Table 1*).

#### G12 mPAE does not alter glutamate transmission in the PL

We first examined potential changes to basal glutamate transmission following mPAE by recording in voltage-clamp configuration. In males we found no effect of exposure on either sEPSCs (frequency: [*t*(14) = 0.82, *p* = 0.425],*Fig. 3B*; amplitude: [*t*(14) = 1.57, *p* = 0.139], *Fig. 3C*) or mEPSCs (frequency: [t(14) = 0.82, p = 0.425], *Fig. 3*C; amplitude: [*t*(14) = 1.17, *p* = 0.262], *Fig. 3D*), indicating that mPAE did not affect AP-dependent or -independent glutamate transmission. Similarly, in females we found no differences in sEPSCs (frequency: [*t*(12) = 1.94, *p* = 0.07], *Fig. 3A*; amplitude: [*t*(12) = 0.15, *p* = 0.167], *Fig. 3B*) or mEPSCs in females(frequency: [*t*(12) = 1.65, *p* = 0.125], *Fig. 3C*; amplitude: [*t*(12) = 1.64, *p* = 0.127], *Fig. 3D*). Together, these results indicate that neither AP-dependent nor independent glutamate transmission onto PL layer 2/3 pyramidal neurons was affected by mPAE.

#### G12 mPAE decreases the excitability of PL pyramidal cells

To determine whether the excitability of PL pyramidal cells was altered by mPAE, we next conducted experiments in zero-current configuration (to determine RMP) and current clamp configuration (for AP firing experiments). We first assessed whether mPAE produced changes to the basal membrane properties of PL pyramidal cells (*Supplementary Table 1*). In males, we did not find any differences on the membrane capacitance [t(13) = 0.33, p = 0.743] or membrane resistance [t(13) = 0.2, p = 0.804]. We also did not find differences in these variables in females [capacitance: t(10) = 0.78), p = 0.455; resistance: t(10) = 1.57, p = 0.147]. When assessing changes to RMP, we found no difference in males [*t*(13) = 0.81, *p* = 0.434] or females [*t*(10) = 0.32, *p* = 0.759](*Supplementary Table 2*). In males, there was also no difference for rheobase, or the amount of current required to fire the first AP [*t*(13) = 0.39, *p* = 0.704]. However, there was a significant difference on rheobase in females [*t*(100 = 3.21, *p* = 0.009]](), indicating that cells from mPAE females required more current to begin firing APs (*Supplementary Table 2*). There was also a significant difference in males on the amount of time required to fire the first AP [*t*(13) = 2.23, *p* = 0.044] demonstrating that cells from mPAE males required more time within a current step to fire the first *AP*(*Supplementary Table 2*). There was no effect of exposure on this variable in females [*t*(10) = 0.29, *p* = 0.778]] (*Supplementary Table 2*). In males, there were no effects of exposure on the peak of the first AP fired [*t*(13) = 0.99, *p* = 0.342] or the first AP threshold [*t*(13) = 0.83, *p* = 0.424] (*Supplementary Table 2*). There was also no effect of exposure in females on either of these variables [peak: *t*(10) = 0.94, *p* = 0.373; threshold: *t*(10) = 0.81, *p* = 0.439] (*Supplementary Table 2*).

When examining the firing patterns of PL pyramidal cells while injecting a series of depolarizing current steps in males, we found a significant effect of exposure [*F*(1, 170 = 4.69, *p* = 0.044] as well as a significant step x exposure interaction [*F*(14, 2265) = 2.32, *p* = 0.005] (*Fig. 4A&C*). In females, we also found a significant main effect of exposure [*F*(1, 100 = 6.78, *p* = 0.026] as well as a step x exposure interaction [*F*(14, 140) = 3.09, *p* = 0.0003](*Fig. 4B&D*). These firing results indicate that, while all cells increase the number of APs fired per depolarization step, cells from mPAE offspring fired fewer at each step than their air-exposed counterparts. Additionally, there was a significant increase in rheobase in cells from mPAE females, indicating that these cells also require more stimulation to begin firing. Overall, our assessment of the excitability of PL pyramidal cells following mPAE demonstrate that mPAE decreases the excitability of PL pyramidal cells in both sexes, although potentially through different mechanisms.

#### Inhibitory modulation of PL pyramidal cell excitability is disrupted following G12 mPAE

Finally, to determine whether mPAE may have affected inhibitory modulation of PL pyramidal cell firing, in a separate subset of cells we examined whether applying gabazine (10 μM) would differentially affect firing in air-vs. mPAE-exposed offspring. When analyzing the effect of gabazine on the firing properties in males, we found no effect of exposure on rheobase [*t*(13) = 0.023, *p* = 0.982], or RMP [*t*(13) = 0.51, *p* = 0.622](*Supplementary Table 3*). To determine whether there were differences in inhibitory modulation of the firing pattern of pyramidal cells, we selectively compared the number of evoked APs between air- and mPAE-exposed cells in the presence of gabazine. This analysis revealed no effects of exposure, with no main effect of exposure [*F*(1, 13) = 0.004, *p* = 0.951] and no current injection x exposure interaction [*F*(14, 182) = 0.18, *p* = 0.999](*Fig. 4E&G*). When analyzing the firing properties of the first AP, there was also no difference in the time to first AP [*t*(13) = 1.04, *p* = 0.319], first AP peak [*t*(13) = 0.12, *p* = 0.906], or first AP threshold [*t*(13) = 0.73, *p* = 0.481] (*Supplementary Table 3*). Together, these results indicate that gabazine application eliminated preexisting differences in action potential firing between cells from air- and mPAE-exposed offspring.

We conducted the same gabazine experiment in a separate set of cells from female offspring, again to determine whether mPAE may have altered the way GABAergic inhibition modulates pyramidal cell firing. In females, we found no main effect of exposure on our measures of rheobase [*F*(1, 16) = 0.03, *p* = 0.959] or RMP [*F*(1, 16) = 0.02, *p* = 0.900](*Supplementary Table 3*). Like in males, we also examined the effect of gabazine on the firing patterns of female pyramidal cells in the presence of gabazine to determine whether inhibitory modulation was different between exposure groups. We again found no main effect of exposure [*F*(1, 16) = 0.45, *p* = 0.513], and no current injection x exposure interaction [*F*(14, 224) = 0.542, *p* = 0.906] (*Fig. 4F&H*). There were also no effects of exposure on time to first AP [*t*(16) = 1.14, *p* = 0.271], or first AP peak [*t*(16) = 1.06, *p* = 0.305]. We did observe a significant exposure effect on the first AP threshold [*t*(16) = 2.91, *p* = 0.010], indicating that cells from mPAE females were more depolarized during gabazine application than air-exposed cells (*Supplementary Table 3*).

Together, these results indicate that gabazine application eliminated preexisting differences in AP firing between cells from air- and mPAE-exposed offspring, suggesting mPAE may increase inhibitory modulation of AP firing in the PL.

## Supplementary Figure Captions

**Supplementary Figure 1.**
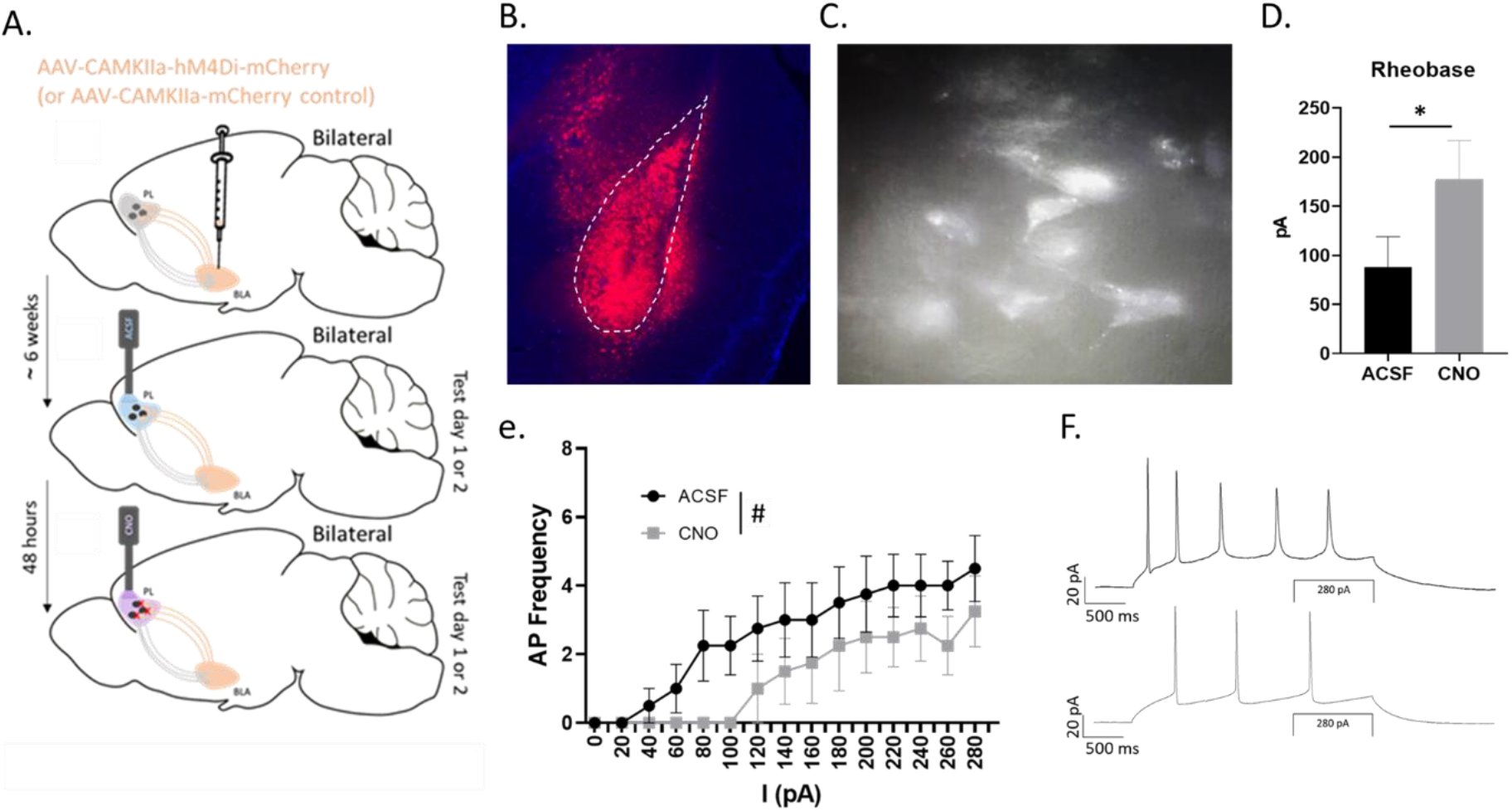
Schematic of chemogenetic experimental design and physiological confirmation of CNO inhibition of inhibitory DREADD virus-transduced cells. **(A)** Schematic showing the experimental procedure and design for the chemogenetic behavioral experiment. Adult offspring were infused bilaterally into the BLA with either an inhibitory DREADD virus (pAAV-CaMKIIa-hM4D(Gi)-mCherry) or a control virus (pAAV-CaMKIIa-mCherry). After six weeks, rats received an injection of either ACSF or CNO into the PL and tested on social behavior. 48 hours later, rats received the opposite injection (CNO or ACSF, respectively) and tested again with a novel partner. **(B)** Representative fluorescent image of virus-expressing pyramidal cells in the BLA (white dashed line delineating BLA border). **(C)** Virus-expressing BLA pyramidal cells identified under an mCherry filter on a slice electrophysiology microscope and used for physiological confirmation of inhibition by CNO. CNO **(D)** increased the rheobase and **(E)** decreased the number of APs elicited by increasing current injections, indicating CNO reduced the excitability of virus-containing pyramidal cells. **(F)** Representative traces of ACSF and CNO recordings. * indicates a significant effect of CNO (*p*<0.05); # indicates a trend (n=4) toward an effect of CNO.

**Supplementary Figure 2.**
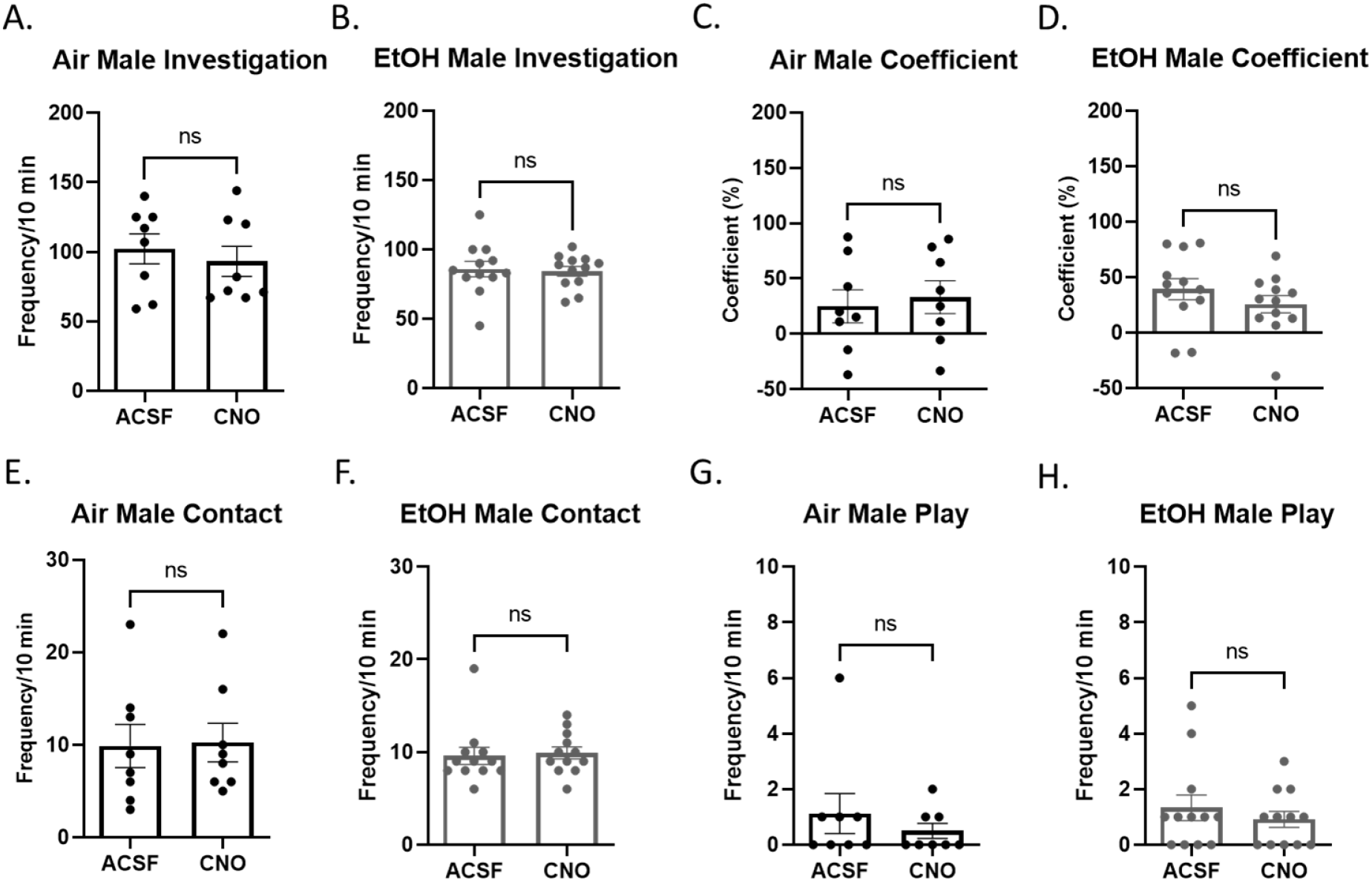
Social behavior following injection of CNO in male rats containing the inactive control virus (pAAV-CaMKIIa-mCherry). There were no effects of CNO injection on any behaviors in air males **(A-D)** or mPAE males **(E-H)**.

**Supplementary Figure 3.**
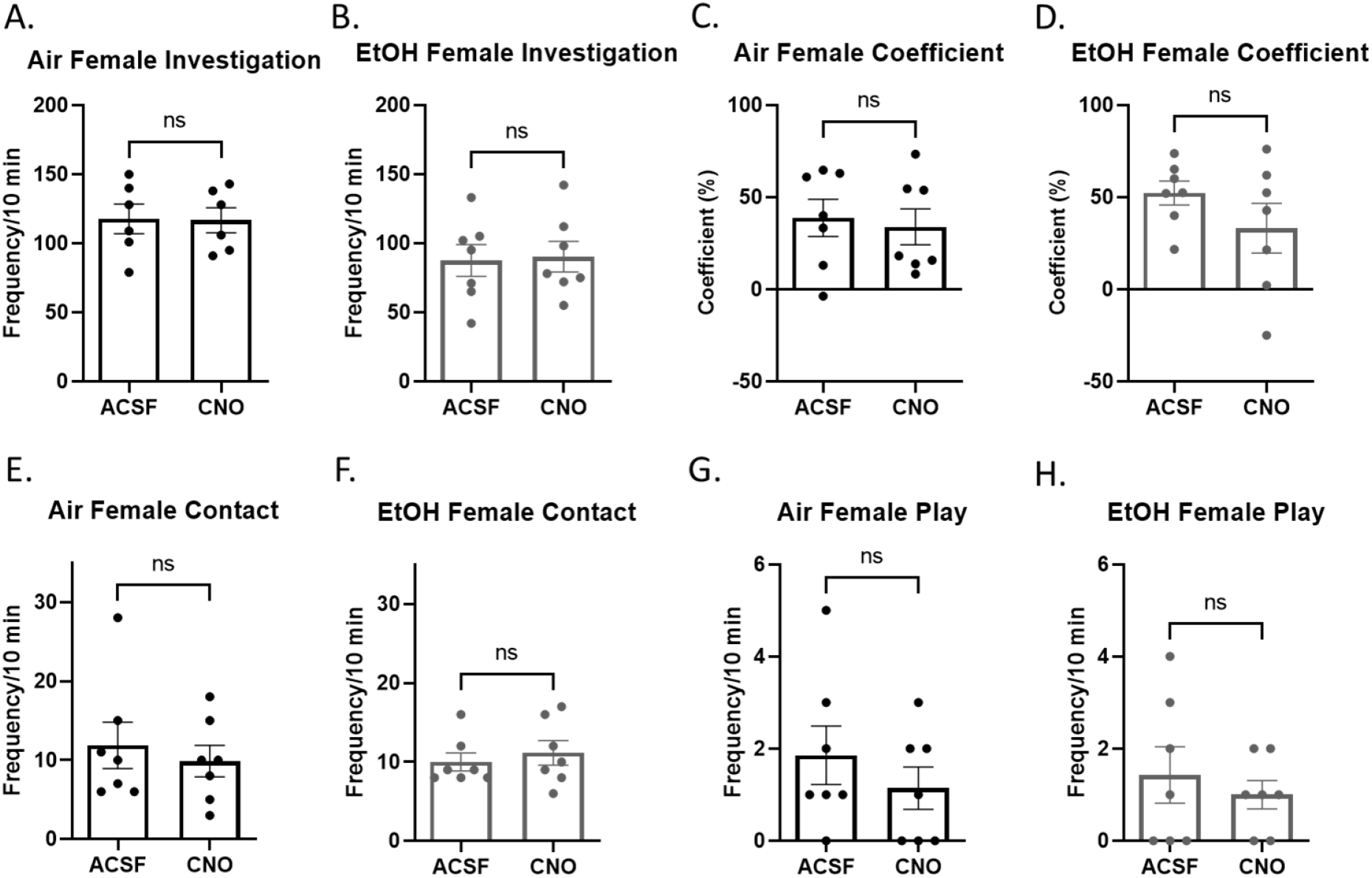
Social behavior following injection of CNO in female rats containing the inactive control virus (pAAV-CaMKIIa-mCherry). There were no effects of CNO injection on any behaviors in air females **(A-D)** or mPAE females **(E-H)**.

## Supplementary Table Captions

**Supplementary Table 1.** Membrane properties of PL layer 2/3 pyramidal cells across exposure and sex, reported as mean (SEM). There were no exposure-induced differences in membrane capacitance or membrane resistance in males or females.

|  | **Air Males**<br>**(n = 6)** | **EtOH Males**<br>**(n = 9)** | **Air Females**<br>**(n = 7)** | **EtOH Females**<br>**(n = 5)** |
| --- | --- | --- | --- | --- |
| **Membrane**<br>**Capacitance**<br>**(pF)** | 245.5 (26.8) | 237.2 (10.04) | 242.3 (14.76) | 258.3 (12.83) |
| **Membrane**<br>**Resistance**<br>**(MΩ)** | 92.95 (10.75) | 89.56 (8.33) | 104.3 (12.09) | 80.7 (47.66) |

**Supplementary Table 2.**
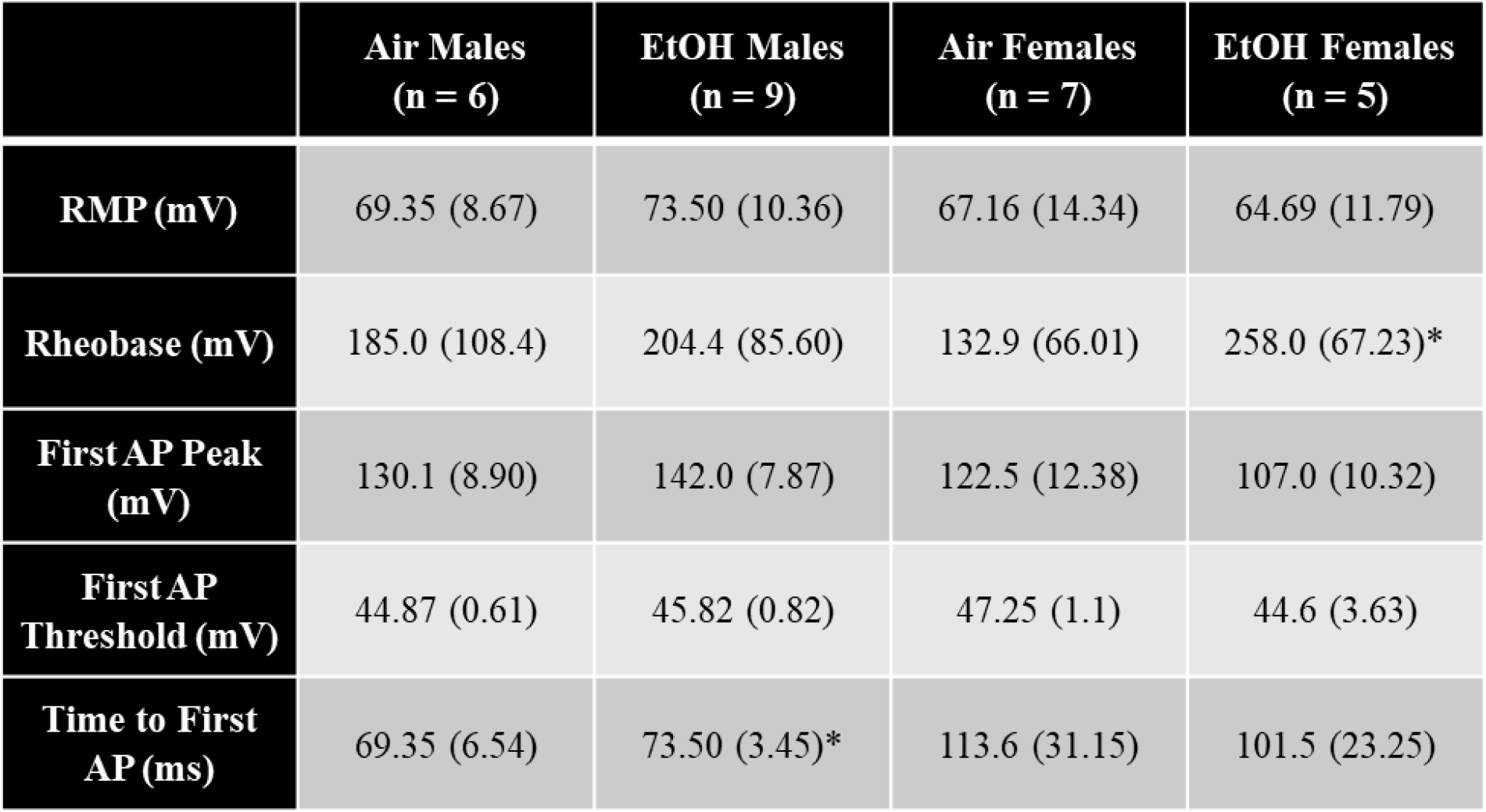
Basal excitability data from PL layer 2/3 pyramidal cells across exposure and sex, reported as mean (SEM). There were no effects of exposure on RMP, first AP peak, or first AP threshold. Cells from mPAE females had increased rheobase compared to control females. Cells from mPAE males had a longer latency to fire the first AP compared to control males. * indicates a significant exposure effect within sex.

**Supplementary Table 3.**
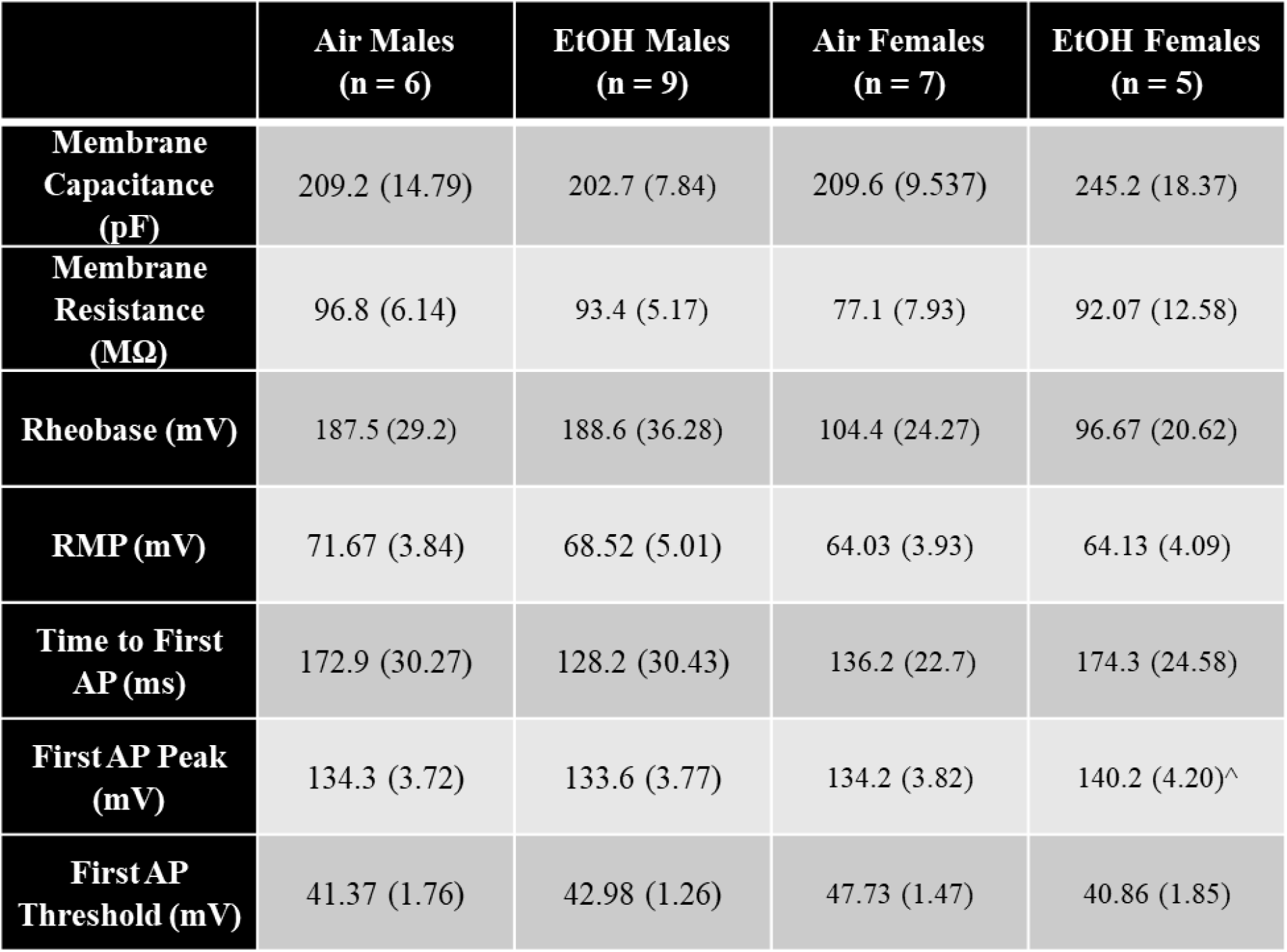
Membrane properties and excitability data from PL layer 2/3 pyramidal cells across exposure) in the gabazine modulation experiment, reported as mean (SEM). There were no differences between exposures on membrane capacitance or membrane resistance on any variable in males. In females, there was a significant increase in the first AP peak of cells from mPAE females compared to air cells. ^ indicates a significant difference between exposure groups within the gabazine application condition (*p*<0.05).

